# Expanding Chemical Space: Developing a Compound Generative Pre-trained Transformer for De Novo Drug Design

**DOI:** 10.1101/2025.01.24.634665

**Authors:** Tze Shin Chen, Jhih Wei Chu, Jinn Moon Yang

## Abstract

The drug development process is time-consuming and costly. With the success of attention mechanism models across various domains, including drug development, their adoption has significantly increased. pre-trained a model on large datasets enhances its ability to understand the vast chemical space, enabling effective drug design. However, despite numerous generative models for drug discovery, challenges remain: (i) Most current models utilize less than 2 million compounds, which are insufficient to cover the vast chemical space and often results in limited interpretability. (ii) Existing pre-trained models for compound generation struggle to represent the full diversity of potential molecular structures due to restricted training data. Here, we developed a generic compound generator leveraging extensive chemical spaces. Training data derived from 200 million compounds in the ZINC20 database enabled the model to capture and represent the SMILES syntax and the key features of compounds. Using the attention mechanisms of the generative model, we achieved an interpretable understanding of chemical structures, ensuring robust performance in diverse chemical spaces. Our approach demonstrated 99% novelty, surpassing state-of-the-art methods to generating chemically valid and unique compounds.

## 1 Introduction

The total number of potential drug-like candidates is estimated to range from 10^23^ to 10^60^ [1], yet only approximately 10^8^ molecules have been synthesized to date [2].

Despite substantial advancements in computational chemistry and bioinformatics over the past decades, efficiently exploring and characterizing this vast landscape of potential compounds remains a critical obstacle in drug development.

Traditional computational methods, such as molecular docking [3], Quantitative Structure-Activity Relationship (QSAR) [4], pharmacophore modeling [5], and unsupervised learning [6], have provided valuable tools for narrowing this space. Recent innovations, including deep learning [7] and Generative Adversarial Networks (GANs) [8], have further expanded the capabilities of computational drug design. Despite these advancements, the complexity of the vast chemical space demands even more innovative approaches.

The emergence of deep learning technologies, particularly attention-based models, has introduced new opportunities in this domain. Autoencoders [9] have enabled data-driven continuous representations of molecules, facilitating the generation of novel compounds. Generative recursive networks [10] have focused on creating specific molecular structures. Since the introduction of the transformer architecture in 2017 [11] attention-based models have achieved groundbreaking success in capturing complex dependencies within sequential data.

In drug discovery, these models have shown exceptional promise. For instance, MolGPT [12] employs a transformer decoder to generate molecular structures conditioned on specific properties. Similarly, DrugGPT [13] integrates protein and ligand inputs, enabling the model to learn intricate relationships between biomolecules. These advances underscore the transformative potential of pre-trained attention-based models in exploring and understanding the vast chemical space, laying the foundation for innovative drug discovery methodologies.

With the availability of extensive data and deeper insights into specific diseases, de novo drug design has made significant strides. Recent deep learning models, particularly those that leverage SMILES, a one-dimensional string representation of molecules, have demonstrated remarkable potential in drug discovery. Models such as Autoencoders, RNNs [10], and GPT [14] have been shown to effectively generate complete molecules. Despite these advancements, several challenges persist:

- **Limited exploration of vast chemical space:** Models trained on datasets like MOSES [15] or GuacaMol [16], which include fewer than 2 million molecules, fail to fully explore the vast chemical space. Furthermore, their “black-box” nature obscures the knowledge they have learned.
- **Under representation of chemical diversity:** Most past approache used single-atom encodings [9] [10] [12] [17], which is simple and direct but fails to capture the broader chemical diversity in molecular structures. This method treats atoms as independent units, ignoring key relationships between functional groups, ring systems, and other molecular moieties.
- **Challenges with attention mechanisms:** While attention-based models are powerful, their application in molecular generation often lacks systematic interpretability, hindering their integration into chemical design processes.

To address these issues, this study proposes a pre-trained framework based on large-scale datasets(Zinc20 [18]) called Compound Generative Pre-trained Transformer (CompGPT). By incorporating frequent fragment encoding(FCS) [19] and leveraging Generative Pre-trained Transformers (GPT) combined with attention mechanisms, the framework aims to comprehensively explore chemical space and enhance model interpretability. The goal is to develop a method capable of capturing key features of chemical structures and generating molecules with high novelty and diversity. Additionally, visualizing attention weights can reveal the chemical information learned by the model, addressing limitations in interpretability and chemical diversity in current models.

## Materials and Methods

This section outlines the methodology used in our research, structured into four key components as illustrated in Figure 1 : dataset preparation, molecular representation, model architecture, and training procedure with evaluation metrics. By combining these components, we aim to address the challenges of exploring the vast chemical space efficiently while ensuring the generation of high-quality, diverse, and novel compounds.

**Figure 1.**
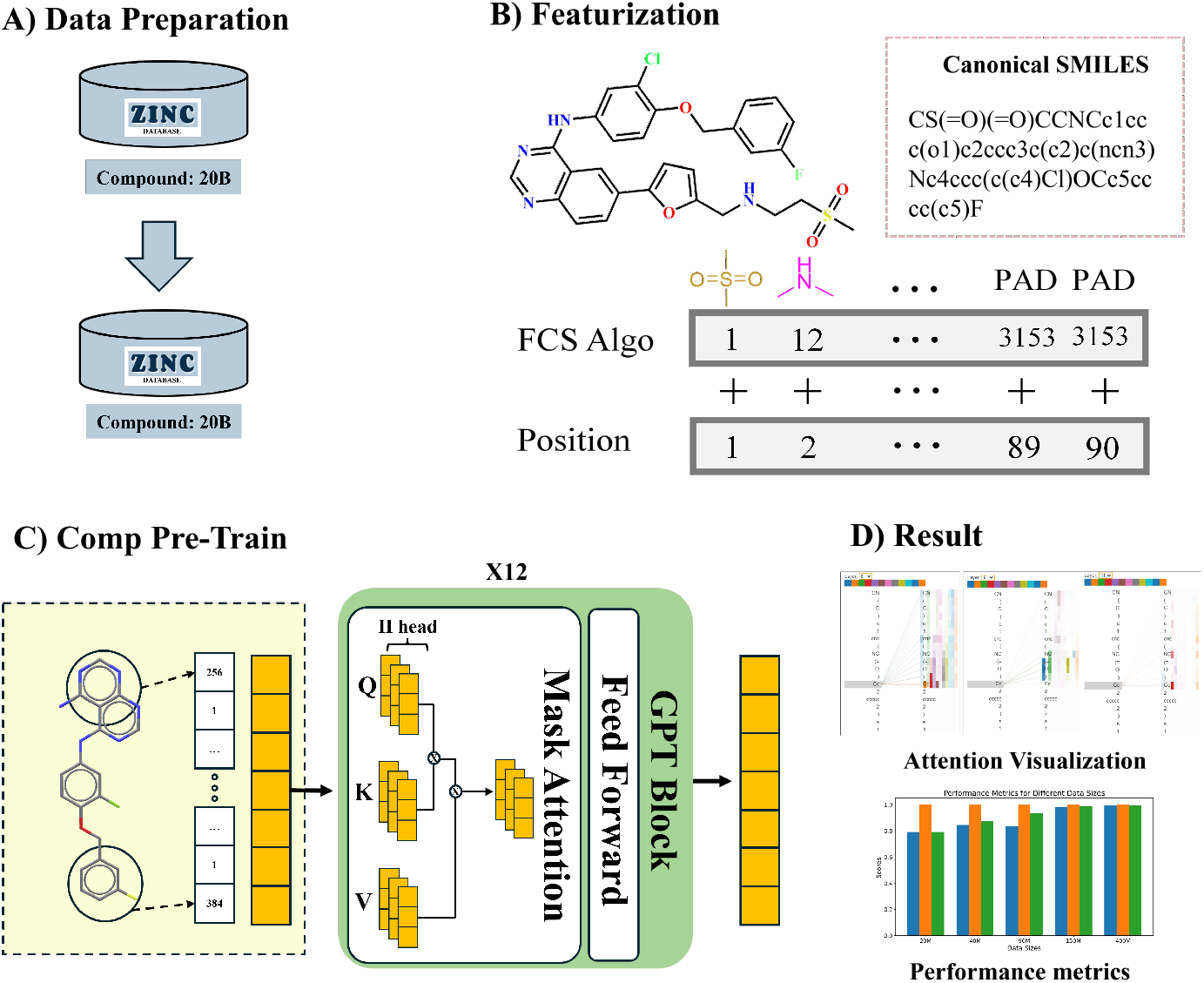
(A) To ensure of large chemical space while maintaining training efficiency, we selected 200M representative ligand compounds from 20B molecules in the ZINC20 for model training. (B) All compounds are processed using the Frequent Consecutive Subsequence (FCS) algorithm to identify and encode frequently occurring fragments as single tokens, facilitating the model’s understanding of meaningful moieties. (C) The model was first pre-trained on a large dataset of molecular SMILES to learn the general patterns and substructures of molecules. (D) CompGPT performance

### Dataset Preparation

ZINC20 is a large-scale public database that aggregates nearly all commercially available compounds, encompassing a vast chemical space with a total of 1.4 billion compounds.

In our research training a general model covering as wide a chemical space as possible, we aim to utilize the entire ZINC20. However, due to hardware limitations and the need for efficiency, we have established a compromise between exploring the extensive chemical space and maintaining manageable data volumes. To determine the data size that yields the most efficient model convergence without sacrificing significant coverage, we have randomly selected a subset of 200 million compounds from ZINC20 for training our general model, ensuring optimal utilization of the chemical space available.

### Molecular representation

Molecular representation is critical for the compound pre-trained GPT and the Ensemble Expert Guide System. In this study, molecules were represented using SMILES [20] (Simplified Molecular Input Line Entry System), a linear notation that translates molecular graphs into character sequences. To enhance the model’s ability to learn and generate chemically valid structures, we employed a tokenizer based on the Frequent Consecutive Subsequence (FCS) algorithm [19]. The FCS process identified 3,151 frequently occurring substructures, such as functional groups and ring systems, from the training dataset. These substructures were tokenized as individual units, reducing sequence complexity while preserving essential chemical information.

Additionally, positional encoding was integrated to capture spatial relationships between substructures, enabling the model to better comprehend molecular arrangements.

### Model architecture

The compound pre-trained GPT is designed to explore the vast chemical space by learning the grammar and structural features of molecular representations. The model architecture is based on GPT-2, which utilizes the transformer decoder. This architecture is particularly suited for sequential data, such as SMILES strings, because of its ability to capture long-range dependencies through the attention mechanism. The generator focuses on generating novel and valid molecular structures by leveraging large-scale training data.

To allow the model to recognize the positional relationships within sequences, positional encodings are added to the embedding layer at the first stage of the transformer decoder. These encodings are calculated using sine and cosine functions:

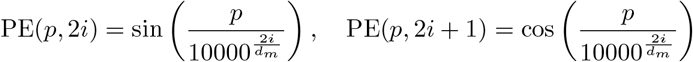

where *p* is the position, *i* is the embedding dimension, and *d*_*m*_ (here set to 768) is the embedding size. These encodings ensure that the model processes tokens in parallel while preserving information about their relative positions in the sequence. This positional information improves the generator’s ability to represent structural relationships within SMILES strings.

At the core of the model is the multi-head self-attention mechanism, which allows it to process relationships across tokens within a sequence effectively. Each decoder module incorporates multiple key components that work together to ensure robust learning. The multi-head self-attention sub-layer identifies dependencies between tokens by focusing on different parts of the input sequence simultaneously, capturing information about functional groups or entire scaffolds of compounds. This is followed by a feed-forward sub-layer.

To further stabilize the training process and ensure smooth convergence, residual connections and layer normalization are applied around each sub layer. The self-attention mechanism itself is mathematically defined as:

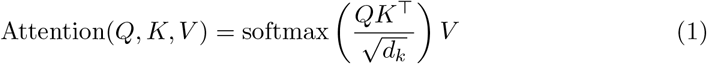

where *Q, K*, and *V* are the query, key, and value vectors, and *d*_*k*_ is the dimensionality of the key vectors. Multi-head attention extends this concept by computing attention across multiple subspaces in parallel, aggregating information from diverse perspectives. The model employs 12 attention heads, with each head having *d*_*k*_ = *d*_*v*_ = 64, balancing computational efficiency and representation diversity.

**Training Procedure and Evaluation Metrics**

Each model was trained for 10 epochs using the Adam optimizer [21] with a learning rate of 2 × 10^−5^. During the generation process, a start token was provided to the network, and the temperature was set to 1. Most models converged and showed optimal performance after 10 epochs. Model performance was evaluated using the following metrics: minimum likelihood (loss), validity, uniqueness, novelty, and internal diversity.

- **Validity (***V* ): The percentage of the generated molecules that conform to chemical rules.

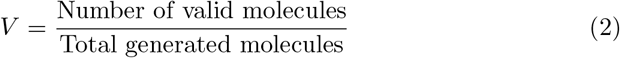
- **Novelty (***N* ): The proportion of generated molecules not present in the training set.

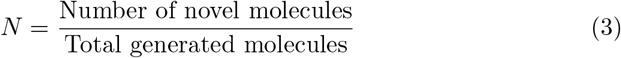
- **Uniqueness (***U* ): The diversity of generated molecules, measured as the ratio of unique molecules to the total generated.

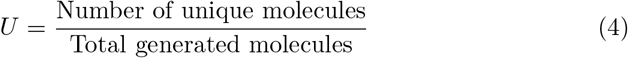
- **Internal Diversity (IntDiv1, IntDiv2)**: Measures the structural diversity within the generated set. It is computed as the average pairwise Tanimoto similarity between molecular fingerprints, subtracted from 1.

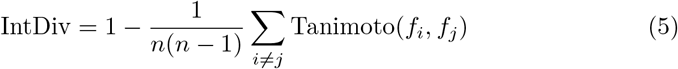

## Results

In this section, we divide into three parts for validation. The first part evaluates the compound pre-trained GPT, including a comparison of its internal parameters and different encoding methods, And the second part, we assess its performance against existing state-of-the-art methods. The last part explores the interpretability of the attention, examining how the model captures key features of molecular structures to ensure that its generated outputs are reasonable and maintain diversity.

### Performance Comparison with Data size and Encoding methods

#### Data size

The ZINC20 contains over 2 billion compounds, making it essential to determine the optimal dataset size for model convergence. To address this, we pre-trained a GPT-2 model on datasets of varying sizes: 20M, 40M, 90M, 150M, and 400M compounds, each trained for 10 epochs.

To assess the impact of dataset size on model convergence, we analyzed the training loss. As shown in Figure 2, the model exhibited significantly better convergence when the dataset size exceeded 150M compounds compared to smaller datasets. The loss values stabilized and reached lower levels, indicating improved learning capacity and generalization. However, increasing the dataset size beyond 150M compounds resulted in only marginal improvements, suggesting diminishing returns in convergence performance. Table 1 summarizes the results across different dataset sizes. The Uniqueness metric remained consistently high, demonstrating the model’s robust ability to generate diverse compounds. Meanwhile, the Validity and Novelty metrics improved substantially with larger dataset sizes, peaking at 150M compounds. Beyond this threshold, further gains in performance plateaued, indicating minimal benefits from larger datasets.

**Table 1.** Model performance metrics based on data sizes Compares GPT-2 performance across different data sizes. Validity and novelty improve with larger datasets, peaking at 150 million data points, while uniqueness remains consistently high (0.999).

| Data size / GPT-2 | Validity (%) | Uniqueness (%) | Novelty (%) |
| --- | --- | --- | --- |
| 20 million | 78.7 | 99.9 | 78.7 |
| 40 million | 84.3 | 99.9 | 87.4 |
| 90 million | 83.2 | 99.9 | 93.0 |
| 150 million | 97.8 | 99.9 | 98.3 |
| 400 million | 99.2 | 99.9 | 99.0 |

**Figure 2.**
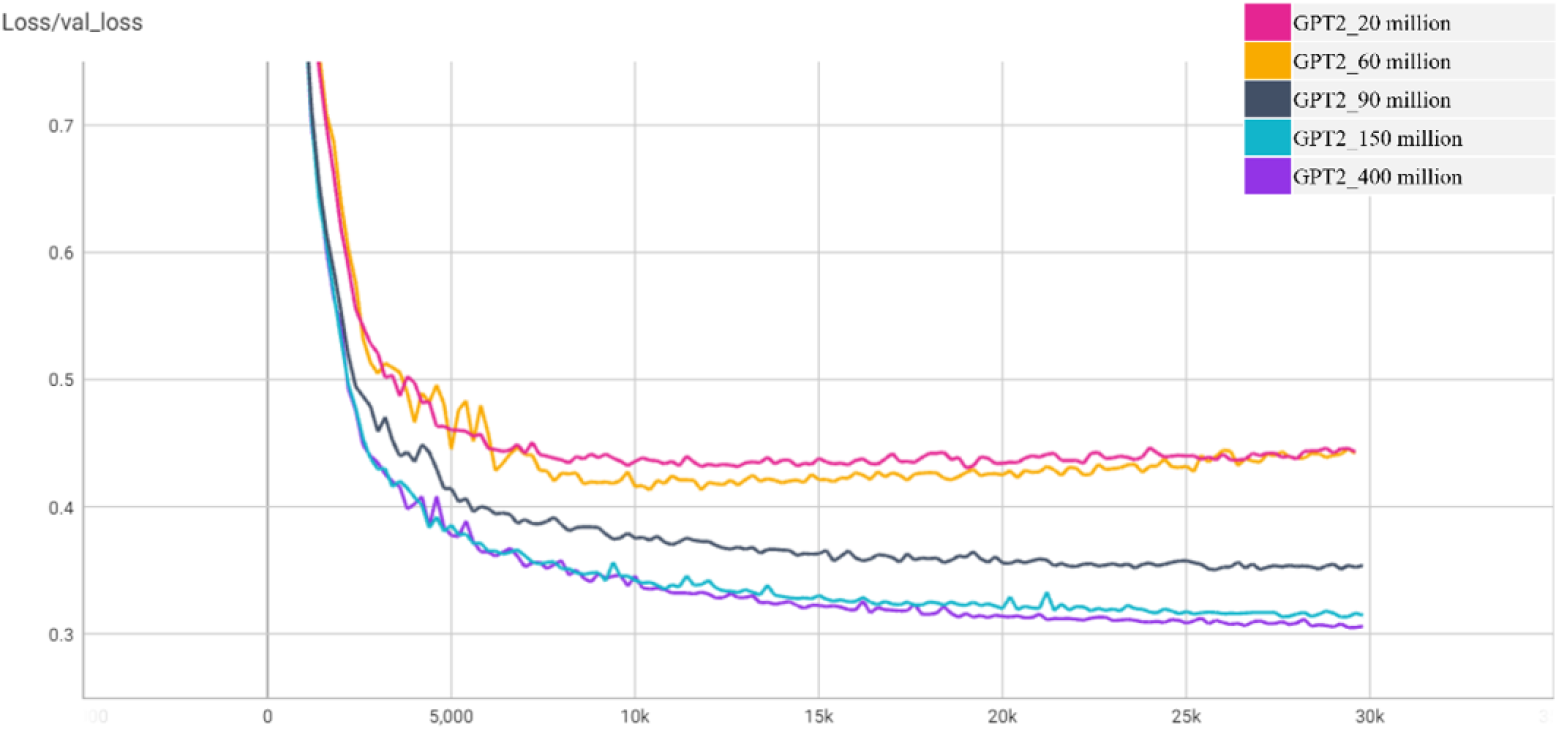
Training loss between different dataset sizes and visualize by Tensor Board [22].

Training with approximately 150M compounds strikes an optimal balance between computational efficiency and chemical space coverage, enabling the generation of high-quality, diverse, and novel compounds. These findings indicate that a dataset size between 150M and 200M compounds achieves the best trade-off between performance and efficiency. Based on this conclusion, we selected 200M compounds for training to ensure comprehensive chemical space coverage. Additional analyses of the extracted samples and their relationship to the original data are presented in Supplementary Material S1.

#### Encoding methods

To further enhance model performance, we compared two encoding methods: atom-level encoding (one-hot), which represents molecules as individual atomic units, and moiety-based encoding, which utilizes FCS to encode meaningful molecular substructures as single tokens. As shown in Table 2, both approaches demonstrated high Validity and Uniqueness. However, moiety-based encoding significantly outperformed atom-level encoding in Novelty and Internal Diversity (IntDiv1 and IntDiv2). This underscores the importance of encoding molecular substructures to improve the model’s ability to generalize and generate diverse compounds. By capturing essential chemical patterns, moiety-based encoding facilitates the exploration of a broader chemical space.

**Table 2.** Performance Comparison of Different Encoding Methods in CompGPT. Compares the performance of CompGPT (Atom) and CompGPT (Moiety) in terms of validity, novelty, and internal diversity. The results show that CompGPT (Moiety) outperforms CompGPT (Atom) across all metrics, with particularly notable performance in validity (99.6%) and novelty (99.9%), as well as higher diversity.

| Encoding Method | Validity (%) | Uniqueness (%) | Novelty (%) | IntDiv1 | IntDiv2 |
| --- | --- | --- | --- | --- | --- |
| CompGPT (Atom) | 92.3 | 99.9 | 93.0 | 0.856 | 0.832 |
| CompGPT (Moiety) | **99.6** | **100** | **99.9** | **0.881** | **0.872** |

### Model Performance Comparison

The chemical space is practically infinite and largely unexplored, making it essential for generative models to produce a significant number of novel and valid molecules to facilitate effective exploration. High scores in metrics such as Validity, Uniqueness, and Novelty ensure that models effectively learn molecular grammar without overfitting, while Internal Diversity (IntDiv1, IntDiv2) indicates the extent of chemical space traversal.

We compared CompGPT with benchmarked approaches, including CharRNN [23], VAE [24], AAE [25], MolGPT [12], LatentGAN [26], and JT-VAE [27], using the MOSES dataset and a generation temperature of 1.0 (10,000 generated SMILES), as summarized in **Table 3**. Among these, JT-VAE employs graph-based inputs, whereas the others use SMILES representations.

**Table 3.** Comparison of benchmarked approaches based on Validity, Uniqueness, Novelty, and Internal Diversity (IntDiv1 and IntDiv2).

| Model | Validity (%) | Uniqueness (%) | Novelty (%) | IntDiv1 | IntDiv2 |
| --- | --- | --- | --- | --- | --- |
| CharRNN | 85.2 | 96.0 | 94.2 | 0.85 | 0.72 |
| VAE | 87.7 | 98.1 | 95.1 | 0.86 | 0.73 |
| AAE | 89.5 | 98.2 | 94.8 | 0.86 | 0.74 |
| MolGPT | 93.0 | 98.9 | 96.5 | 0.87 | 0.76 |
| LatentGAN | 86.3 | 97.2 | 93.0 | 0.84 | 0.70 |
| JT-VAE | **100.0** | 98.0 | 97.5 | 0.88 | 0.75 |
| **CompGPT** | 99.5 | **99.2** | **98.8** | **0.89** | **0.77** |

JT-VAE achieves perfect Validity by verifying molecular correctness at every generation step. Excluding JT-VAE, our model achieves nearly perfect scores in Validity (99.5%) and Uniqueness (99.2%), demonstrating a robust ability to learn SMILES grammar through its attention mechanism, which effectively handles long-range molecular dependencies. Furthermore, our model outperforms all the others in Novelty (98.8%), showcasing its capacity to explore new chemical regions, thanks to its extensive training dataset and larger model parameters.

In terms of Internal Diversity, our model exhibits slightly greater chemical diversity compared to other methods, with IntDiv1 and IntDiv2 values of 0.89 and 0.77, respectively. This highlights its ability to generate chemically meaningful and diverse structures while matching or exceeding the performance of existing approaches. These results collectively demonstrate the effectiveness of our approach in producing valid, novel, and diverse compounds, while maintaining a balance between exploration and specificity within the chemical space.

To further illustrate the model’s capabilities, we analyzed the structural similarity between model-generated compounds and known kinase inhibitors, such as EGFR and FGFR3 inhibitors. For EGFR, the model consistently captured key moieties that aligned closely with critical binding residues like Met793 in docking simulations. Similarly, for FGFR3, the model successfully generated compounds with moieties highly consistent with known inhibitors, ensuring strong interaction with FGFR3’s binding pocket. These examples underline the model’s capacity to generate highly relevant compounds tailored to specific kinase targets, adapting effectively to different chemical profiles over training epochs.

JT-VAE achieves perfect Validity by verifying molecular correctness at every generation step. Excluding JT-VAE, our model achieves nearly perfect scores in Validity and Uniqueness, demonstrating a robust ability to learn SMILES grammar through its attention mechanism, which effectively handles long-range molecular dependencies.

Moreover, our model outperforms all others in Novelty, showcasing its capacity to explore new chemical regions, thanks to its extensive training dataset and larger model parameters. In terms of Internal Diversity, our model exhibits slightly greater chemical diversity compared to other methods, underscoring its ability to generate chemically meaningful and diverse structures while matching or exceeding the performance of existing approaches.

### Explanation of the Compound Pre-trained GPT

In GPT-based models, the transformer decoder architecture is utilized, with the self-attention mechanism playing a crucial role in capturing long-range chemical structure relationships within compounds. By leveraging interactions between Query and Key (K), the model determines the relevance of each component in predicting the next structure. To enhance interpretability, we visualize these attention transformations and verify whether the model has effectively learned critical information about the compounds. As shown in Figure 3,additional data and analyses are provided in Supplementary Material S2. A visualization of attention weights across 12 layers and 12 heads highlights how different layers and heads learn distinct levels of information. In the initial layers, the attention mechanism focuses on global relationships, identifying the relative position of the next element within the compound. In the middle layers, attention shifts to local structural features, such as aromatic rings or functional groups. Finally, in the last layers, attention is focused on understanding how different moieties within the compound are connected, revealing a learned comprehension of the scaffold and the relationships between substructures. This attention behavior demonstrates the model’s progressive refinement of its chemical knowledge. The early layers provide a global perspective, enabling the model to contextualize molecular elements, while the subsequent layers dive into localized and detailed structural relationships critical for generating valid and coherent chemical structures. These visualizations validate that the self-attention mechanism is instrumental in determining which chemical structural information to prioritize during the generation process.

**Figure 3.**
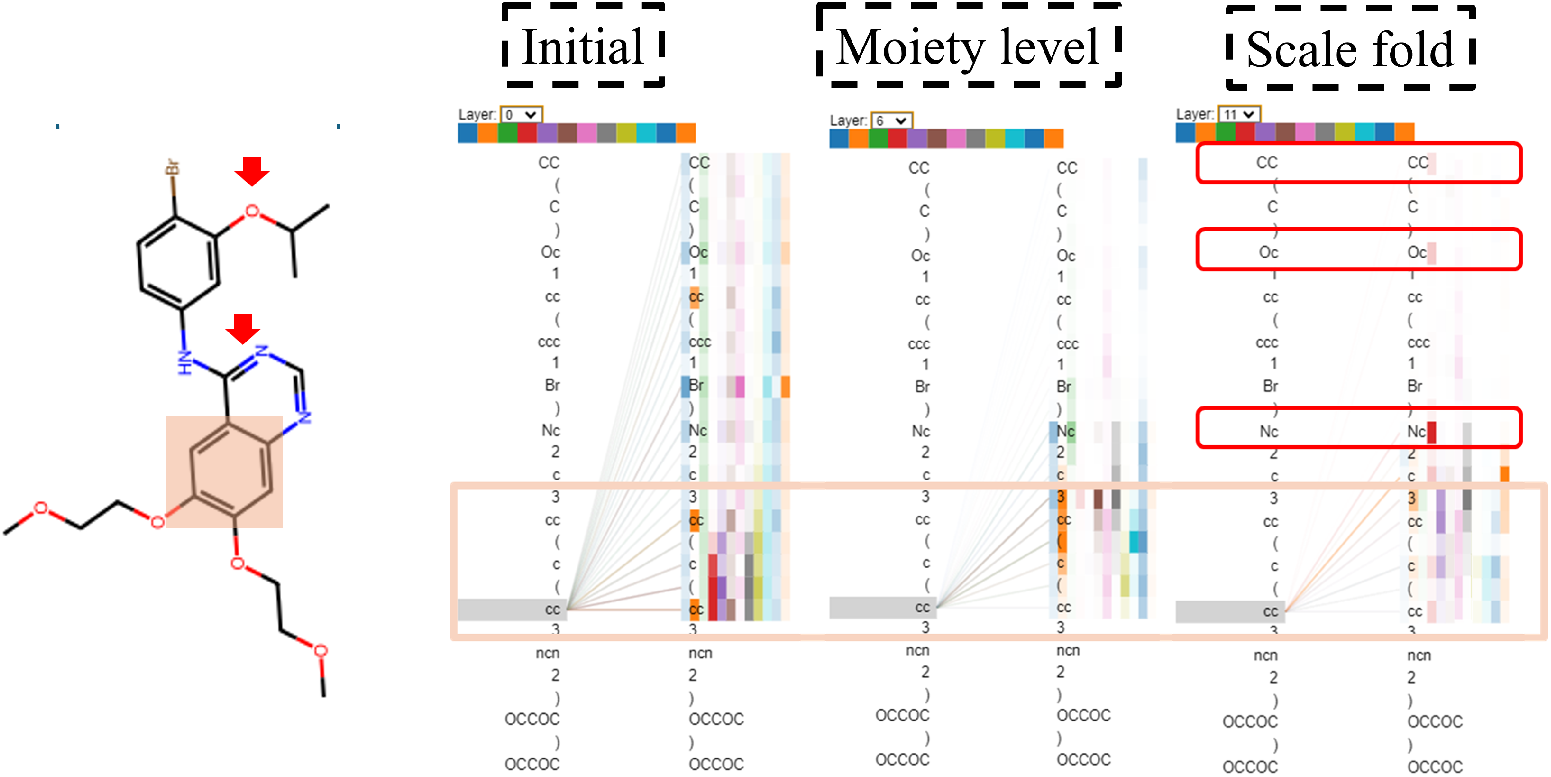
Attention weight visualization showing atomic, moiety-level, and Scale fold importance in chemical structures, with key features highlighted.

## Discussion

This study demonstrates the potential of leveraging large-scale datasets and advanced attention-based generative models for drug discovery. By pre-training on 200 million compounds from the ZINC20, our approach addresses challenges such as insufficient dataset sizes, limited chemical diversity, and a lack of interpretability in existing models. The use of FCS encoding enables the generation of diverse and chemically meaningful compounds by capturing important molecular substructures. Training with 200M compounds strikes an optimal balance between computational efficiency and chemical space coverage, while moiety-based encoding significantly improves novelty and internal diversity compared to atom-level encoding. Visualizing attention weights reveals how the model progressively refines its understanding of molecular structures, from global relationships to functional groups and moiety connectivity, ensuring generated compounds are logical and diverse. Compared to state-of-the-art models, our approach achieves superior novelty and internal diversity while matching the validity of graph-based methods like JT-VAE. This work establishes a robust framework for generative drug discovery, combining large-scale data, advanced encoding techniques, and interpretable attention mechanisms, with opportunities for future extensions in scalability and target-specific applications.

## Code availability

## Supporting Information

**Figure S1.**
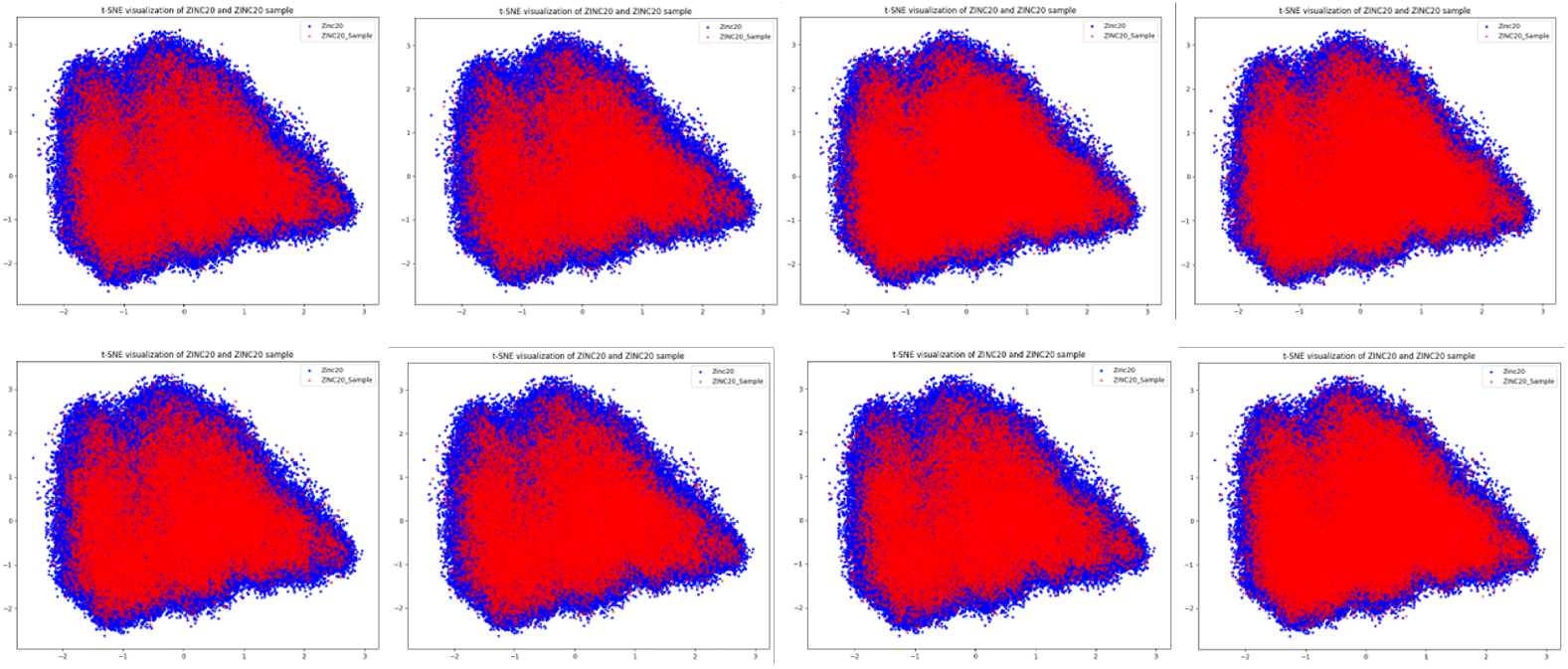
The t-SNE visualization compares the chemical space distribution of the entire ZINC20 (blue) and the randomly sampled subset (red). The analysis demonstrates that the subset effectively captures the primary chemical space characteristics of the ZINC20.

**Table S1.**
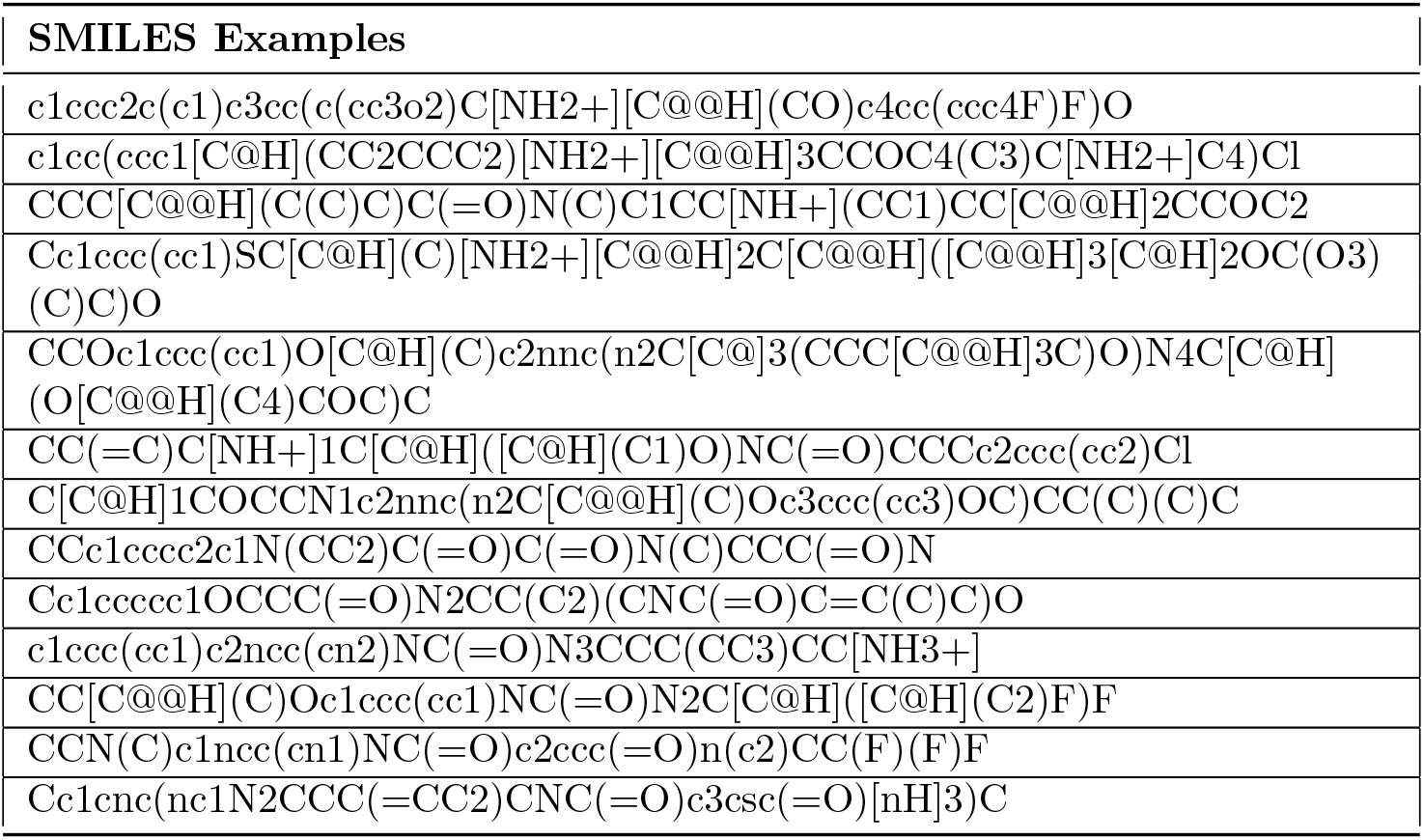
Examples of SMILES generated by fine-tuned CompGPT. The table includes randomly selected samples to demonstrate the model’s ability to produce chemically diverse and structurally valid compounds.

**Figure S2.**
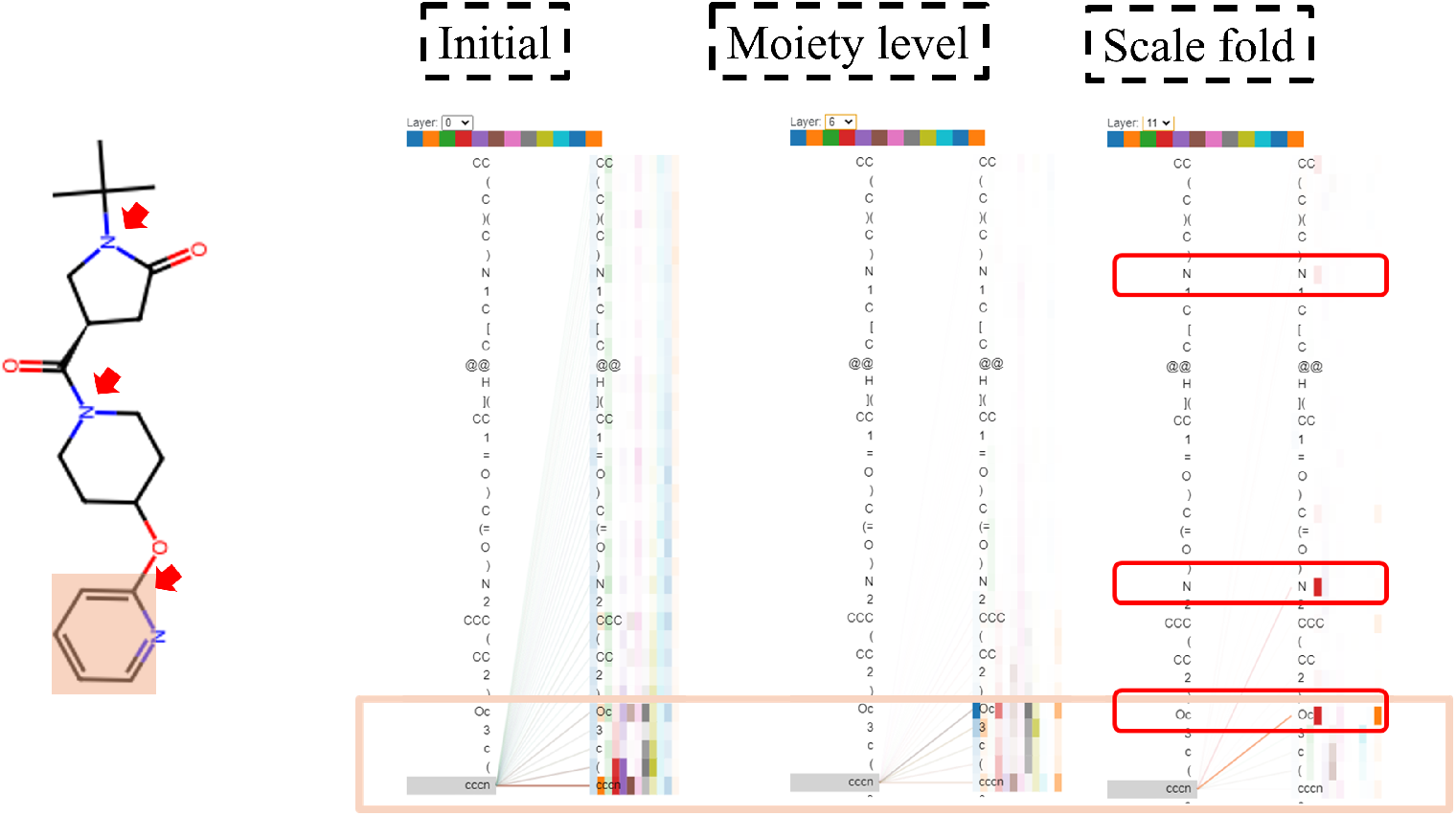
Attention visualization 2

